# Machine learning analysis of lung adenocarcinoma and squamous cell carcinoma microbiome datasets reveals biomarkers for early diagnosis

**DOI:** 10.1101/2023.11.25.568645

**Authors:** Pragya Kashyap, Kalbhavi Vadhi Raj, Naveen Dutt, Pankaj Yadav

## Abstract

**Introduction:** Lung cancer (LC) is the second most frequent cancer worldwide with high incidences and mortality rates and non-small-cell-lung cancer (NSCLC) accounts for 80-85% cases of LC. It is further majorly sub-classified into adenocarcinoma (AC), and squamous-cell carcinoma (SCC). A late diagnosis at an advanced stage, a high rate of metastasis, and the development of therapy resistance are responsible for approximately 95% of mortality. Owing to high heterogeneity and variances in subtypes, it is important to precisely classify them for treatment. However, it poses a challenge in clinical practices, as it requires accurate quantification of each proportion subtype which is time-consuming and sometimes erroneous. The lower airways are home to a dynamic bacterial population sustained by the immigration, elimination, and migration of microbes from the gastrointestinal tract and upper airway tracts. The disruption in the homeostasis of microbiome compositions was found to be correlated with the increased risk of LC. Artificial intelligence (AI) techniques are used extensively in the early screening, and treatment of NSCLC have made significant strides in recent years. Recently, the use of CT/MRI scan image data in prediction models also results in false-positive rates and requires subsequent tests for further exploration which delays the prognostication. Therefore, early diagnosis, prevention, and treatment are critical to enhance survival and reduce death. Here, we aim for the classification of AC and SCC using the lung microbiome of lung tissue samples, implementing AI-based algorithms.

**Methods:** We have obtained raw sequencing data from the NCBI online database, and 149 AC samples and 145 SCC samples in patients were analyzed for their lung microbiome present in the lung tissue samples. The metadata such as patient age, sex, smoking history, and environmental material (malignant or not) were also analyzed. Using these data, machine learning algorithms were applied to select the best microbiome features for the classification of subtypes.

**Results:** A supervised ML and DL based model was developed that can discriminate NSCLC subtypes based on their microbial information, exploring the microbiome as predictive information for early screening. Consequently, 17 features were identified as a biomarker, and they showed good performance in distinguishing AC from SCC with an accuracy of 81% in KNN and 71% in DNN when demonstrated on the validation dataset.

**Conclusion:** This study proposed a supervised machine learning framework where we can rely on taxonomic features and AI techniques to classify overlapped AC and SCC metagenomic data providing lung microbiome as a predictive and diagnostic biomarker in LC. Moreover, our framework will also be very helpful to other researchers to obtain further biomarkers and perform analysis in overlapped subtypes in different diseases.

## Introduction

Lung cancer (LC) is the second most prevalent cancer throughout the world, with high incidences and mortality rates. As per GLOBOCAN 2020, approximately 1.79 million people die from LC annually and non-small-cell-lung cancer (NSCLC) accounts for 80-85% cases of LC [1]. NSCLC, are further majorly sub-classified into adenocarcinoma (AC), and squamous-cell carcinoma (SCC). A late diagnosis at an advanced stage, a high rate of metastasis, and the development of therapy resistance are responsible for approximately 95% of mortality [2]. Despite advancements in therapeutic strategies, NSCLC patient survival has not increased significantly. In the United States, the relative survival of 5-year for patients with stage I was 65 %, while the patients with stage IV were less than 10 % [3]. Even if the patient underwent surgical resection, the post-operative recurrence rate was still high. In fact, 30-55% patients develop postoperative recurrence despite complete curative resection [4]. Targeted and contemporary immune-based therapy have minimally improved NSCLC survival[5]–[7]. Implementing preventive strategies complemented with early screening and efficient therapeutics can lower the worldwide NSCLC prevalence, incidence, and mortalities.

The recent advancements in next-generation sequencing technologies and analytical methods have provided new information on the lung microbiome, which was earlier thought to be sterile, for its potential in clinical interventions. Growing research has suggested that an imbalanced lung microbiota, termed “dysbiosis” influences can cause carcinogenesis through several different possible mechanisms [8]–[10]. These mechanisms include microbiome dysbiosis, genomic instability, affecting metabolism, induction of inflammatory pathways, and immune response in the host. For example, studies have shown that certain bacteriotoxins and other pro-inflammatory factors from microorganisms such as *Haemophilus influenzae*, E. coli, Enterobacter spp., Moraxella, and Legionella genera are correlated that drive the inflammation of lung tissue and contribute to the formation of tumours, to promote carcinogenesis [11]. Liang et al., indicated that *Mycobacterium tuberculosis* (TB) may cause LC through chronic inflammation-associated carcinogenesis. This suggests the causative relationship between specific microbes and the development of cancer [12].

Recently, studies also highlighted that machine learning (ML) and deep learning (DL) algorithms are useful for LC-associated lung microbiota characterization and prediction [13], [14]. Moreover, most of the research used microbial communities to differentiate cancer from healthy controls [15]. However, whether lung tissue microbial profiles can distinguish NSCLC subtypes for early identification complemented with ML and DL based technologies is unknown. Both AC and SCC have significant heterogeneity and different features, prognostic consequences, and therapeutic responses. Due to the variations in treatment approaches for different NSCLC subtypes, their proper classification is of utmost importance in clinical settings [16], [17]. Therefore, recent research focuses on AI-based technologies for the diagnosis and treatment of NSCLC. Teramoto et al., trained the CNN model with an accuracy of 71.1% by analysis of 76 cytological images of NSCLC types revealing that the results obtained were consistent with the classification conclusions provided by cytopathologists [18]. Khosravi et al., distinguished between AC and SCC using fine-tuned Convolution Neural network (CNN) model in localized highly magnified images with an accuracy ranging from 75% to 90% [19]. However, a lot of Magnetic Resonance Imaging (MRI) / Computed Tomography (CT) scan images or image data result in false positives, which prompt several tests and biopsies that each have their own morbidities [20]. Despite research on these subtypes, the classification of subtypes in the early stages of cancer continues to pose a challenge for medical professionals. This challenge arises from the lack of consensus among pathologists and the occurrence of multiple morphological subtypes within one or more sections of the same patient. In order to achieve an accurate diagnosis, it is imperative to quantify each subtype, in a process that is both time-consuming and prone to inaccuracies [21].

Emerging evidence, however, suggests that most patients exhibit overlapping lung microbiome signatures in NSCLC subtypes with varying compositions [22]. This complicates classification with traditional ML and DL learning techniques, which deserve further exploration. Hence, it is of the utmost importance to identify the biomarkers precisely and promptly in an early stage of NSCLC and modify the treatment regimes for different stages. Therefore, accurate and effective biomarkers are critical for patients with NSCLC for early screening and better survival prognosis. In this work, an attempt has been made to develop a diagnostic model. For this purpose, we used 149 and 145 lung tissue samples of AC and SCC patients, respectively. Samples were collected from the NCBI database. Additionally, data on patient age, sex, smoking history, and environmental material (malignant or not) was also considered for analysis. These variables together were not analyzed earlier but are important in predicting the NSCLC subtype and whether it is curable or not. We fitted a supervised ML and DL model to identify NSCLC subtypes using microbiological data. This study aims to (1) compare tumoural tissue microbiome composition between NSCLC subtypes, and (2) identify microbiome markers to predict AC and SCC subtypes. The outcome of this work is expected to advance our understanding of AC and SCC.

## Methodology

The entire work plan followed in this study is outlined in **Figure 1** and described in detail below. Summarily, there are several steps from the dataset acquisition from NCBI to the model building using lung microbiome.

**Figure 1:**
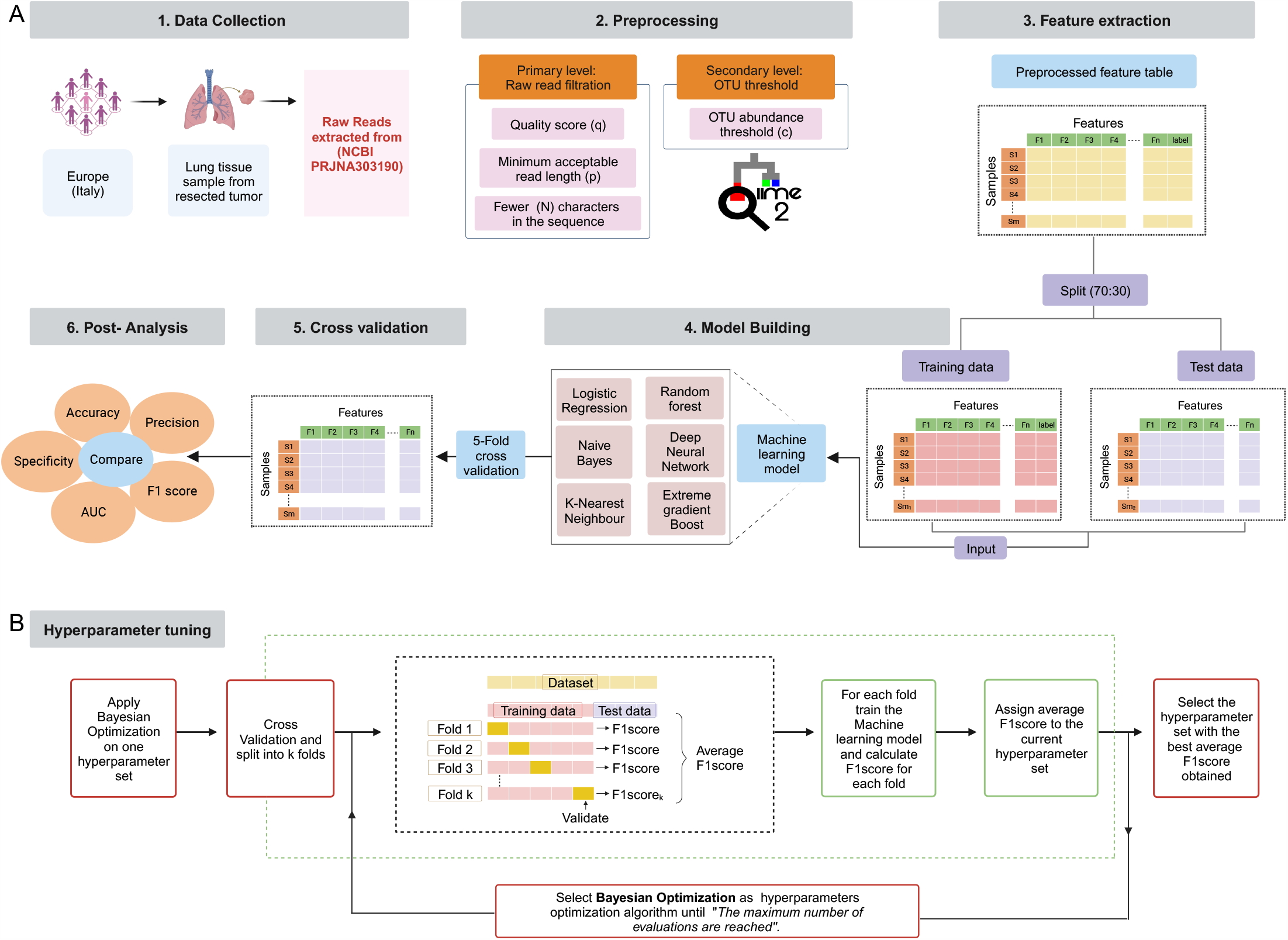
The workflow of the proposed methodology (A) the pipeline followed; (B) shows hyperparameter tuning using the Bayesian Optimization algorithm.

**Figure 2:**
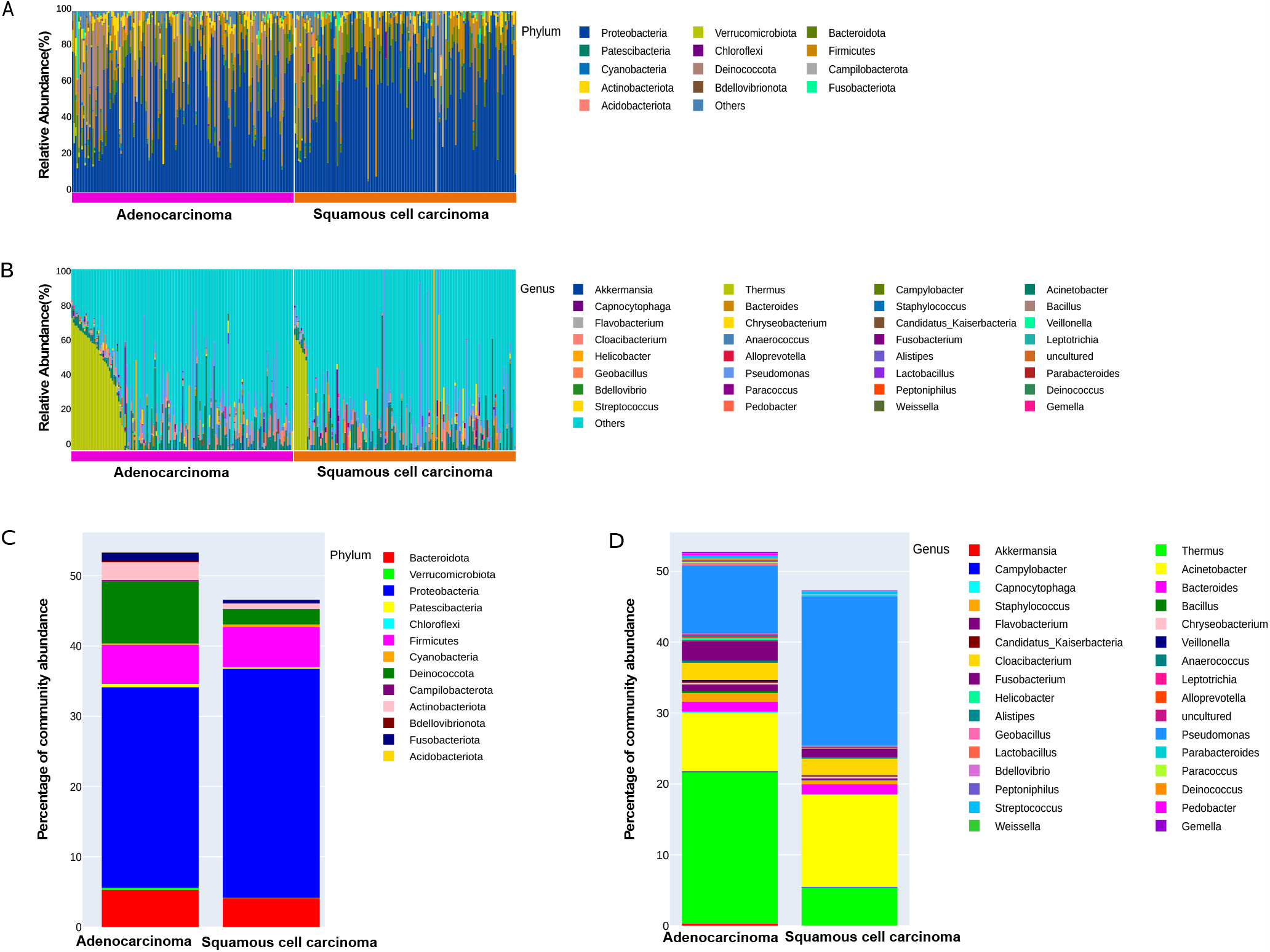
Taxonomic characterization of the lung microbiota in LC patients with (A) AC at Phylum level; (B) SCC at Phylum level; (C) AC at Genus level; (D) SCC at Genus level. Comparison of relative taxa abundance of the lung microbiota in patients with AC and SCC at (A) Phylum level; (B) Genus level. (Only the relative abundance within each group >0.05% are shown)

### 1. Retrieval of data

Raw sequencing data on NSCLC patients was downloaded from NCBI Sequence Read Archive (SRA) using the SRA toolkit (V.2.9.1) with the identifier PRJNA30319 for Yu et al.,[23]. This study used Illumina high-throughput sequencing to target the V3-V4 region of the 16S rRNA gene.

Note: Stage information of patients was retrieved after connecting with the published paper author. Majority of the patients are in the early stage. Only 10 patients were in stage IV, 7 in IIIB and rest were in early stage.

*R/Python codes are submitted to GitHub* (https://github.com/kashpk/Lung-microbiome-biomarker-analysis), (https://github.com/ihdavjar/Lung-microbiome-biomarker-analysis)

### 2. Sequence read processing and taxonomy profiling

The 16S rRNA gene sequencing data was processed using a standardized pipeline implemented in QIIME2 [24]. Briefly, paired-end demultiplexed reads were joined using fastq-pair [25], followed by primary and secondary level filtering as described by Bokulich et al.,[26]. First, the reads were quality filtered based on Phred score (Q<25). Then, reads were checked for ambiguous base calls and read lengths. The reads shorter than 100 bp after trimming were removed. Next, the Deblur approach was used to obtain error-free biological sequences in samples, also known as Amplicon Sequence Variant (ASV) [27], [28]. After quality filtering and denoising techniques, denoised data was then summarized. Consequently, an ASV table resulting in 283 samples was created with a median of 1468 reads/sample (mean of 2429 reads/sample). Representative sequences of each ASV were assessed for their taxonomy lineage employing a sklearn classifier pre-trained on the Silva (v.138-99) using 99% similarity [29]. The sequence reads originating from chloroplast and human mitochondrial DNA were excluded during the Deblur process when they did not match the bacterial 16S database.

Then, sample diversity metrices for α-diversity were assessed using Richness and Shannon diversity. This was done to reflect the microbial diversity within a single sample. The β-diversity was assessed based on Bray-Curtis dissimilarity and visualized using principal coordinate analysis (PCoA) [30]. Afterwards, linear discriminant analysis (LDA) effect size (LEfSe) [31] was used to identify differential taxa abundance among AC and SCC groups.

### 3. Pathway construction

Phylogenetic Investigation of Communities by Reconstruction of Unobserved States2 (PICRUSt2) [32] was used to predict the functional composition of the taxa in AC and SCC samples. Then, the predicted Kyoto Encyclopedia of Genes and Genomes (KEGG) Ortholog (KO) [33] was analyzed descriptively using ggpicrust2 [34] in R.

### 4. Biomarker identification and ML and DL classification models

The LEfSe method was employed to identify distinguishing features between AC and SCC as well as to eliminate redundant features from our dataset. It used the non-parametric Wilcoxon rank-sum test and Kruskal-Wallis to analyze the differences between the two groups. The selected features that had effect sizes greater than 0.5 and threshold of p-value<0.05 were considered statistically significant. However, based on the earlier selected features, certain similarities were observed, particularly between the family Thermaceae and the genus Thermus. Therefore, to mitigate this issue, we conducted a feature selection process by eliminating features that exhibited a correlation coefficient exceeding 0.95 while retaining only one such feature. The outcome yielded a total of 13 distinct features, which were taxa with classification, phylum, class, order, family, genus, and species. Apart from this, four metadata were also included, namely, patient age, sex, smoking status, and environment material (malignant or non-malignant).

We further implemented classification algorithms to determine the predictive potential of certain features selected from LEfSe for analyzing the subtype of NSCLC. The patients of AC were designated as “0”, while the patients of SCC were designated as “1”. Therefore, this problem can be regarded as a binary categorization task. This study employs six classification methods, specifically logistic regression (LR), random forest (RF), extreme gradient boost (XG Boost), naïve Bayes (NB), K-nearest neighbours (KNN), deep neural network (DNN) and modelled in Python 3.6.7 using its standard publicly available libraries. We computed accuracy, precision, specificity, F1 score, and area under the curve (AUC) to evaluate the ML and DL techniques that will help us determine the early-stage subtype. After scoring these parameters, the optimum technique for further analysis was reported.

#### 3.1 Hyperparameter tuning

Supervised ML algorithms are comprised of two categories of parameters: those that undergo optimization during the training process and those that must be specified by the user before initiating the learning process to enhance performance or generalization on the provided dataset. The latter category, known as hyperparameters, is adjusted based on the model’s performance on a validation dataset and is of utmost importance in any ML problem [35]. Conversely, model parameters are determined through data training. Model parameters, such as weights and coefficients, are derived from data by the algorithm.

We used Bayesian optimisation-based hyperparameter optimisation [36]. It is an optimization algorithm that aims to model the loss on the validation dataset as a function of the hyperparameters utilizing a Gaussian Process (GP) and tries to minimize the loss with respect to the hyperparameters. It is considered better than regular extensive search algorithms like random search [37] and grid search [38]. It is computationally more efficient and give global optima, while other optimization algorithms tend to get stuck at local minima. In addition, Bayesian Optimisation offers the benefit of not sampling every search space combination like Grid Search while also being more systematic.

Therefore, in our approach, we divided the dataset, comprising 274 patient samples after the preprocessing steps, into training and test datasets in the ratio 7:3. The division of the datasets is done in such a way that the class ratio is maintained in both datasets. A larger dataset containing 191 samples was used for training, and another dataset containing 83 samples was used for testing. While performing the hyperparameter tuning, training data was subjected to 5-fold cross-validation, and the hyperparameter that gave the best result in 5-fold cross-validation was taken for further studies. Then the ML and DL models were trained with the obtained hyperparameters using training data and finally evaluated using testing data, as shown in **Figure 1C**.

#### 3.2 ML and DL algorithms and their tunable hyperparameters

##### 3.2.1 Logistic Regression

LR is a common classification algorithm. This algorithm aims to fit the sigmoid function to the data in such a way that the loss is minimized. LR has only one hyperparameter, and this is the regularisation parameter, usually denoted by λ. In our study, we are only considering L2 regularisation.

##### 3.2.2 Naïve Bayes classifier

The NB algorithm is a probabilistic and generative classification technique. It assumes that the characteristics are independently distributed and uses the dataset to compute the class-conditional distributions, evidence, and priors. Then the Bayes theorem is used to compute the posterior probabilities. Typically, class-conditional distributions are approximated to Gaussian distributions using the dataset’s mean and standard deviation. There are no adjustable hyperparameters in NB [39], [40].

##### 3.2.3 Random Forest Classifier

An RF classifier is an example of ensemble learning. In this algorithm, multiple decision trees are trained, and the final answer is either the average of all the trees’ outputs or the result of a vote. In this study, we considered the number of trees, the depth of each tree, which is the distance of the lowest node from the tree’s root, the maximum leaf nodes, which is the decision tree’s total leaf nodes, and the minimum samples in a node for further splitting [41].

##### 3.2.4 Extreme gradient boost classifier

XG Boost is also an example of ensemble learning. But unlike the RF algorithm, here each new decision tree is trained on a modified dataset, which is the original training dataset but modified in such a way that the samples misclassified by the previous decision tree are given more weightage compared to the correctly classified data; hence, this method tries to solve the problem of classification by taking microsteps with each new decision tree [42].

##### 3.2.5 k-nearest neighbour

K-nearest neighbour is a non-parametric distance-based machine learning algorithm [43]. In this algorithm, a sample is labelled **L** if k or more samples in its vicinity belong to class **L**. Before training, one of the hyperparameters that must be provided to the algorithm is k. In addition, this algorithm’s distance metric is also a hyperparameter. However, we have not taken this into account and are employing Euclidean distance.

##### 3.2.6 Deep Neural Network

DNNs are a type of neural network with more than one hidden layer. It has wide applications in various fields like computer vision, Natural language processing, etc. However, there is very little research on the application of DNN in biomarker analysis. DNN performance depends on the architecture, layer count, and number of neurons in each layer. Besides layers and neurons, hyperparameters include activation function, regularisation parameter, epochs, batch size, learning rate, and learning rate type (fixed or variable) [44]. Variable learning rates drop steadily as the optimizer approaches the optimal point.

Neural networks link inputs X to responses Y using numerous organized networks of interconnected nodes (neurons) with weights at each edge. Each neuron in feed-forward propagation networks receives input from previous neurons. Backpropagation links input layers (microbial taxonomic feature set; X1, X2,…, Xi) to each neuron in one or more hidden layers to maximize weights and predictive power. Iteratively, an output layer meets the last concealed layer to forecast response output (Y). Neural networks are dynamic and can reveal subtle structures in high-dimensional and complex datasets, making them suitable for studying microorganisms in challenging environments [45].

#### 3.3 Class Overlap Problem

Class overlap occurs when samples from multiple classes share a common region in the data space. These samples share feature values that tend to overlap despite being in different classes or are extremely close to one another, complicating classification with traditional ML and DL techniques [46], [47]. In an ideal classification, the data should be organized so that the inter-class distances are greater than the intra-class distances. However, this is not the case for the majority of real-world problems. Due to the fact that AC and SCC are both subtypes of cancerous cells, the two classes overlap. In our dataset, the distances between classes are not as great as they should be. Consequently, our ML and DL models produced poor results. To address this issue, we have chosen to transform our data with LDA before implementing the previously mentioned ML and DL models.

##### 3.3.1 Transforming the dataset using LDA

LDA is a supervised technique used for dimensionality reduction. The main aim of LDA is to reduce the dimensionality of the dataset while also maximizing the inter-class distances and minimizing the intra-class distances. The dimensionality of the dataset after LDA transformation is less than or equal to one less than the number of classes. In our case, we have only two classes, i.e., AC and SCC. Hence, our resulting dataset contains only one feature. Applying LDA to both the training and testing datasets would add high bias to our study as LDA is a supervised dimensionality reduction method, and in the case of new data, we do not have the labels, as predicting the labels is our problem statement. Hence, applying LDA to training and testing data separately would not be meaningful. Therefore, we are applying LDA to the training dataset. Then we are using the same transformation matrix obtained on the training dataset to transform the test dataset [48].

##### 3.3.2 Model Evaluation and Comparison

The prediction performance of the model in the classification of two groups (AC *vs*. SCC) based on our selected features, we modelled LR, RF, NB, XG Boost, KNN and DNN. All the models were optimized using Bayesian optimisation using 5-fold cross-validation on the training dataset to give the maximum F_1_ score. The evaluation metrics such as sensitivity, specificity, precision, F_1_ score, accuracy, ROC curve, and finally, area under the curve (AUC) are used in this study to evaluate the classification potential of the chosen taxa using the previously mentioned procedure. The metric values are obtained by utilizing a confusion matrix for the purpose of comparing the models. The components of a confusion matrix are depicted in Table 1, and their description is as follows:

**Table1:**
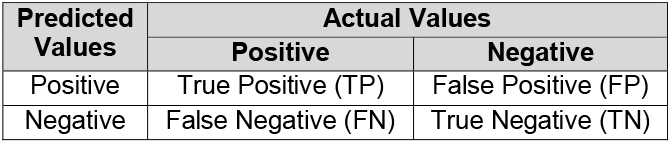
Illustration of confusion matrix.

| Predicted Values | Actual Values |  |
| --- | --- | --- |
|  | Positive | Negative |
| Positive | True Positive (TP) | False Positive (FP) |
| Negative | False Negative (FN) | True Negative (TN) |

**Table1:** Performance Matrix.

| Classifier →<br>↓ Parameters | TN | FN | FP | TP | Accuracy | Precision | Specificity | F1 Score | AUROC |
| --- | --- | --- | --- | --- | --- | --- | --- | --- | --- |
| LR | 33 | 11 | 10 | 29 | 0.75 | 0.74 | 0.77 | 0.73 | 0.85 |
| NB | 32 | 10 | 11 | 30 | 0.75 | 0.73 | 0.74 | 0.74 | 0.85 |
| RF | 36 | 10 | 07 | 30 | 0.80 | 0.81 | 0.84 | 0.78 | 0.86 |
| XGB | 33 | 09 | 09 | 31 | 0.78 | 0.78 | 0.79 | 0.78 | 0.84 |
| K-NN | 36 | **09** | **07** | 31 | **0.81** | 0.82 | 0.84 | 0.79 | 0.84 |
| DNN | 30 | 10 | 13 | 30 | 0.72 | 0.70 | 0.70 | 0.72 | 0.85 |

1. True Positive: TP represents that while finding the type of LC, if the actual value is AC/SCC and the predicted value is also AC/SCC. Then, TP is equal to 1.
2. False Positive If the actual value is AC/SCC, but the predicted value contradicts. Then, FP is equal to 0.
3. True Negative: If the actual is not AC/SCC, but the predicted value is also not AC/SCC. Then, TN is equal to 1.
4. False Negative: If the actual value is not AC/SCC, but the predicted value is AC/SCC. Then, FN is equal to 0.

The evaluation metrics are calculated using the above values:

1. **Accuracy:** It is the most common segmentation and classification metric, defined by the ratio of correct predictions to the total number of predictions as shown in Eq. (1),

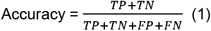
2. **Precision:** It is the ratio of correctly categorized positive samples to total positive samples (either correctly or incorrectly). It is shown in Eq. (2),

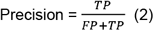
3. **Sensitivity:** It is the ratio of correctly classified positive samples as positive to the number of positive samples. It measures the ability of the model to detect positive samples. It is called recall, shown in Eq. (3),

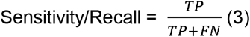
4. **F**_**1**_**-score:** It is the weighted harmonic mean of precision and recall. The best score is 1, and its value is a single number between 0 and 1, derived as shown in Eq. (4),

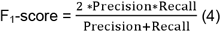
5. **Specificity:** It is the proportion of true negative predictions that are correctly identified. It is shown in Eq. (5).

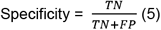

#### 4.5. Statistical Analyses

All statistical analyses have been performed using R 4.3.1 and its required packages. PCoA was carried out utilizing the R package ‘ape’ based on the Bray-Curtis distance matrix. Permutational multivariate analysis of variance (PERMANOVA) [49] with Adonis2 function to evaluate statistical significance between AC and SCC using 10,000 permutations to investigate pairwise comparisons of the groups using the vegan package. This was to check whether overall bacterial composition differed between AC and SCC. The Shannon index and Bray-Curtis dissimilarity, were calculated by using the R package ‘vegan’ and visualized using the ggplot2 package. The LEfSe was used to detect microbial markers and differentiate AC from SCC, with a significance of LDA score > 0.5 and *P*-value < 0.05. Comparison between groups was conducted utilizing the non-parametric Wilcoxon rank-sum test and Kruskal-Wallis.

## 5.4 Results

### 5.4.1 Taxonomic profiling of lung microbiota composition in AC and SCC

We examined AC and SCC microbiota taxon abundance to better understand NSCLC lung microbial community characteristics. The relative taxon abundance in AC and SCC patients on phylum, class, order, family, genus, and species levels were classified and analyzed. The per-sample genus and phylum bacterial taxonomic distribution is shown in **Figure 1** which shows that taxonomic composition varies greatly between individuals. The phylum/genus taxonomic characteristics of the lung tissue microbiota between the AC and SCC groups are presented in **Figures 1A** and **1B**, and the rest of the microbiota taxonomic profiles based on the class, order, and family level are given in **Figure S1**. At the phylum level, Proteobacteria, followed by variation in Bacteroidota, Firmicutes, and Deinococcota were the most common in both the AC and SCC groups **(Figure 1A)**. At the genus level, Thermus, Acinetobacter, Bacteroides, and Staphylococcus were the core genera present in the samples **(Figure1B)**. Additionally, at the family level, the relative frequency of Pseudomonaceae, Comamonadaceae, Moraxellaceae, Thermaceae, and Weeksellaceae was increased more significantly in both groups as shown in **Figure S1**.

**Figure 1C** and **1D** shows the comparison of relative taxa abundance of the lung tissue microbiota in AC and SCC patients at the phylum and genus levels. The microbiome composition which was present in at least 0.05% of samples was shown. AC was found to be dominated by the phyla Proteobacteria followed by Deinococcota and Firmicutes whereas SCC was to be dominated by Proteobacteria followed by Firmicutes and Deinococcota in **Figure 1C**. The genus Thermus dominated the microbiome, followed by Pseudomonas and Acinetobacter in AC whereas Pseudomonas dominated the microbiome, followed by Acinetobacter, and Thermus in SCC **(Figure 1D)**. Additionally, at the family level, Thermaceae dominated the microbiome followed by Comamonadaceae, Moraxellaceae, and Pseudomonadaceae in AC whereas in SCC samples Pseudomonadaceae dominated followed by Moraxellaceae, Comamonadaceae and Weeksellaceae which are displayed in **Figure S2**. Also, at the class level, Gammaproteobacteria dominated the microbiome, followed by variation in Deinococci and Bacteroidia whereas at order level Pseudomonadales dominated, followed by Burkholderiales and Thermales in both groups **Figure S2**.

### 5.4.2 Bacterial diversity and community structure analysis or Microbial diversity and richness and variation in AC & SCC

To measure differences in taxonomic diversity among AC and SCC groups, the richness and Shannon diversity index was calculated. Comparatively, the taxonomic diversity of samples from the AC group and SCC group was significantly distinct. Richness and Shannon index revealed that α-diversity was higher in AC than in the SCC group with a statistically significant p-value of 8.043e-06 and 0.0465 respectively (Fig. 3*a*). To evaluate the significant variation of microbiome structure in AC and SCC, ecologic distances calculated based on the bray-curtis and, were visualized by principal coordinates analysis (PCoA plot) (Fig. 3b). The statistical significance between the groups was tested using PERMANOVA with 10,000 permutations, employing the adonis2 function. The resulting p-value was 3e-04. There was no clear distinction between samples from the AC group and the SCC group. There was relatively weak clustering between the AC and SCC groups.

**Figure 3:**
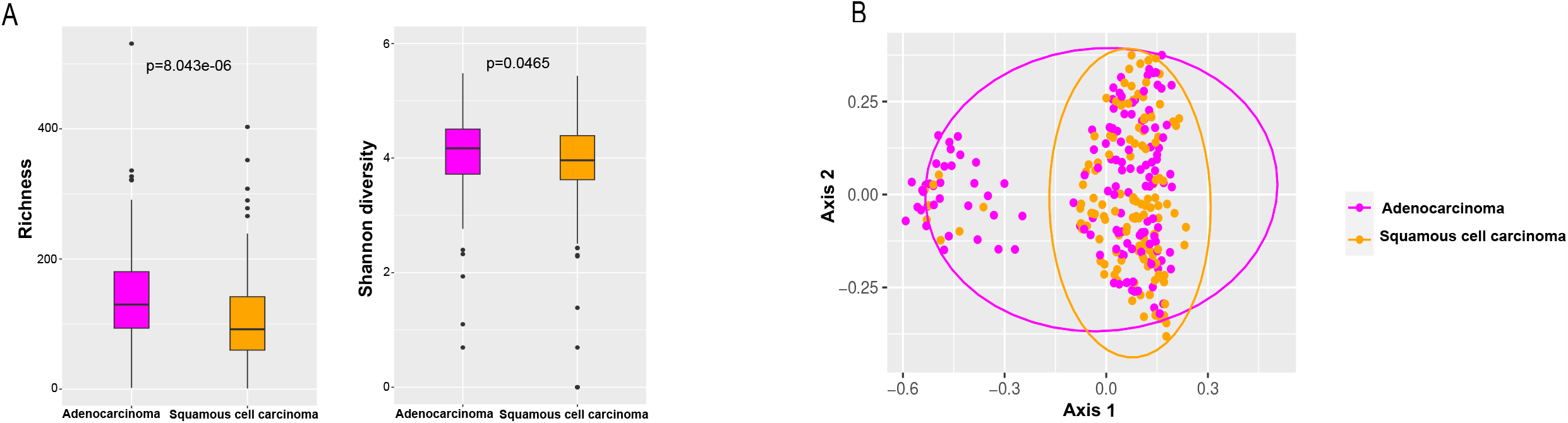
Comparison of α-diversity and β-diversity of the lung microbiota between AC & SCC (A) Shannon, was used to evaluate the α-diversity in AC & SCC. The p-values were computed using the Wilcoxon rank sum test. (B) difference in lung microbiota composition between the two groups was displayed by principal coordinate analysis (PCoA) based on Bray-Curtis metrics, (p-value calculated based on PERMANOVA test). The orange and magenta dots represent SCC and AC, respectively.

### 5.4.3 Analysis of metabolic pathway

The differential functions of bacterial communities were analyzed using PICRUSt2, the predicted KEGG-KO analysis and visualization in the ggpicrust2 package. The differences between the AC and SCC are displayed in **Figure 4**. Overall, 25 were reported to be enriched with adj. p-value less than 0.05. The pathways whose relative abundance is greater than 0.05 were peroxisome, bacterial secretion system, citrate cycle (TCA cycle), RNA polymerase, carbon fixation pathways in prokaryotes, oxidative phosphorylation, Ubiquinone and other terpenoid-quinone biosynthesis, and glutathione metabolism identified in AC and SCC group. Moreover, based on the log2 fold change values, Proteosome and Caffeine metabolism were upregulated, while the Renin-angiotensin system and Hypertrophic cardiomyopathy were downregulated, respectively, despite their low relative abundance, in AC and SCC groups despite their low relative abundance. The microbiota in the two groups always showed obvious metabolic behaviours for the pathways of biosynthesis of other secondary metabolites, xenobiotics degradation and genetic information processing for transcription and folding, sorting, and degradation to be most affected. Further classification of KEGG functional predictions into cellular processes, human diseases, metabolism, organismal systems, and environmental information processing suggested that the different metabolic functions of the microbiota may be an important factor influencing the progression of lung cancer **(see Table S1)**. These data indicated the potentially interesting metabolic and genetic reprogramming in the lung tissue microbiota during lung cancer development.

**Figure 4:**
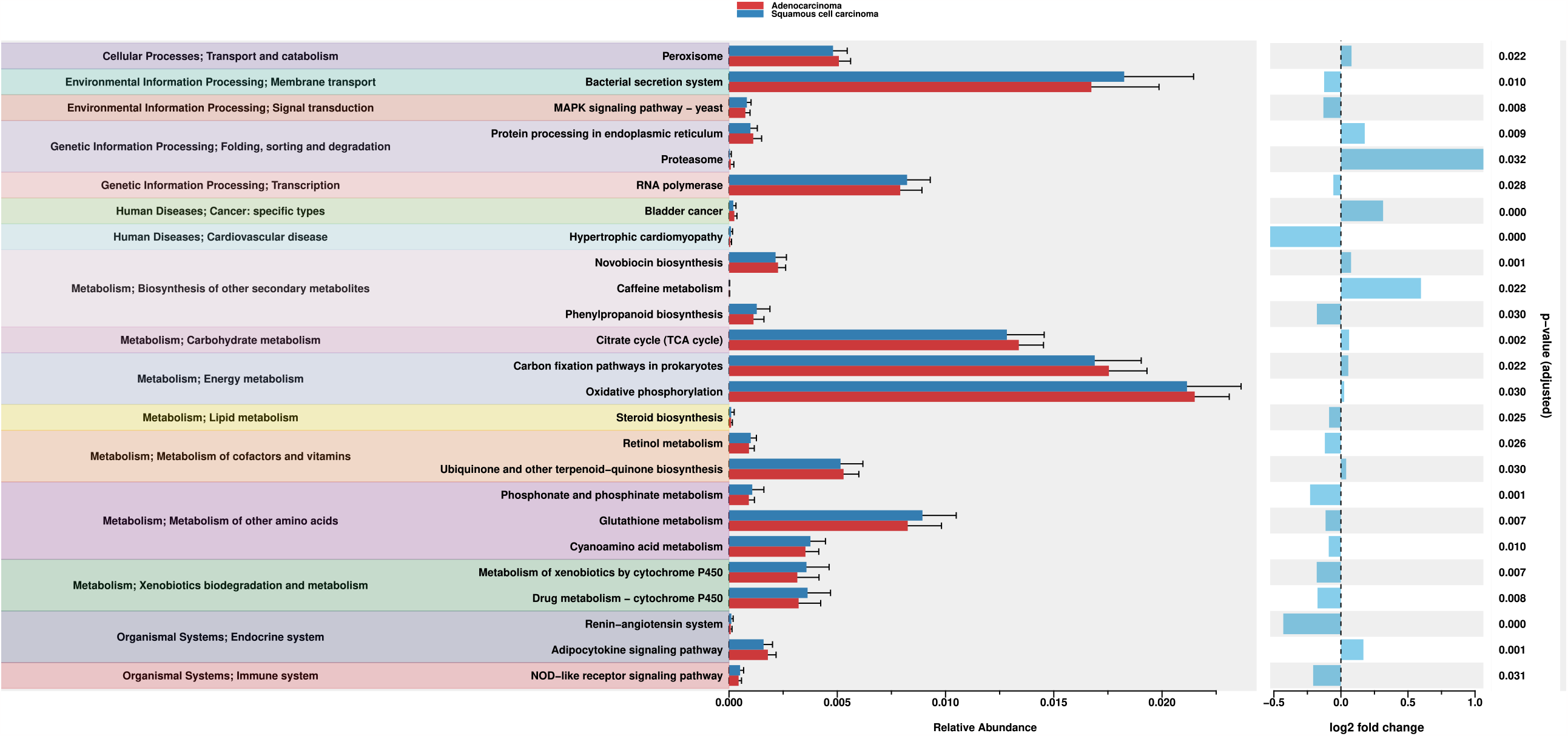
Pathway enrichment analysis in the lung microbiota using PICRUSt2.The impact of differentially enriched KEGG pathways between AC and SCC groups was evaluated through the adj. p-value. Only KEGG pathways meeting an adj. p-value < 0.05 are shown. The columns in red and blue represent the AC and SCC groups, respectively.

### 5.4.4 Differential microbiota compositions in AC & SCC

The LEfSe is an algorithm used for exploring potential bacterial biomarker, by detecting taxa with differential abundance among the two groups (AC vs. SCC). This resulted in identifying the specific lung tissue bacterial taxa associated with two groups. An LDA score above 0.5 indicated the greatest difference in taxa from the phylum to the species level in **Figure 5A**. A total of 359 unique taxa at different levels with significant abundance differences across the two groups were identified. Out of which 32 differentially abundant taxa at different levels were noted, including phylum Deinococcota, order Enterobacteriales, genus Thermus, phylum Actinobacteriota, etc. differentially abundant, in AC. On the other hand, SCC was enriched with the order Psuedomonales, genus Actinobacter, and family Moraxelleace as shown in **Figure 5A**.

**Figure 5:**
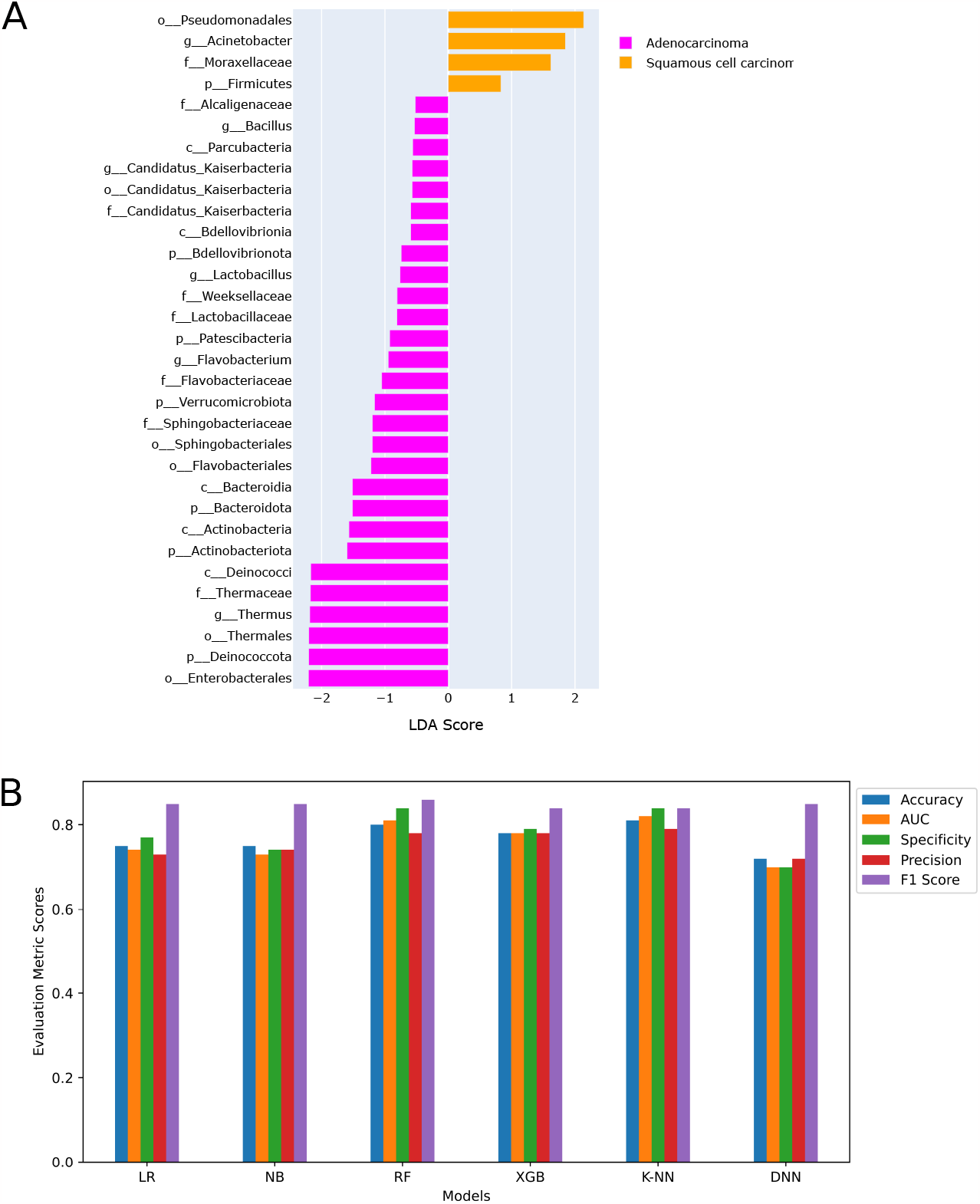
Enriched taxa in AC & SCC lung microbiota are represented at (A) taxa level. (B) Selected features at taxa level showing model performance.

After LEfSe, Pearson correlation was applied to remove the highly correlated features and achieved 13 features for our classifier. For instance, genus Thermus, and order Thermales, belonged to the same classification and had a correlation coefficient greater than 0.95. Therefore, only one feature from those was selected (results of the correlation heatmap are in **Figure S3 and Pearson correlation in Table S2**). Hence, the 13 features after including metadata results in 17 features that were used for model training were family Thermaceae, order Enterobacterales, phylum Actinobacteriota, phylum Bacteroidota, order Flavobacteriales, order Sphingobacteriales, phylum Verrucomicrobiota, family Flavobacteriaceae, phylum Firmicutes, family Moraxellaceae, genus Acinetobacter, order Pseudomonadales, phylum Deinococcota along with environment material (malignant or non-malignant), host sex, host age, and smoking history **(details given in Table S3)**.

### 5.4.5 LDA transformation of dataset

Applying LDA transformation to our dataset reduced its dimensionality while maximizing the separation between AC and SCC. **Figure S4** shows the dataset after LDA transformation. LDA identifies new axes in such a way that the variance of the original dataset is conserved while at the same time maximising the inter-class distance and minimising the intra-class distances. In our case, as the number of classes is two, LDA reduced our number of features to one. The maximum number of features after LDA transformation is always one less than the number of classes present in the dataset. Then, we used our LDA-transformed data for the classification of two classes using the previously mentioned features.

### 5.4.5. Model Evaluation

Given the differences in lung microbiome in AC and SCC, we hypothesized that the selected bacterial features might classify these subtypes along with metadata information in patients with LC. To this end, we constructed several machine learning classifiers and deep neural network to identify AC from SCC using the differential features. The top 17 features after LEfSe and LDA selection were chosen as features for the classification task. **Table 1** shows the performance matrix of classifiers modeled on several parameters such as Accuracy, Precision, Specificity, Sensitivity, and Area Under the Curve (AUC). K-NN achieved the highest accuracy of 81% in accurately classifying the two subtypes followed by RF with 81%. K-NN also showed fewer false positives and negatives compared to other classifiers which is of utmost importance as the correct classification of cancer subtypes determines the further prognosis and treatment strategies.

Additionally, a deep multilayer perceptron was developed, consisting of an input layer including one neuron, one hidden layer containing 217 neurons each, and an output layer with one neuron representing the AC and SCC subtypes. In order to mitigate the issue of overfitting, the implementation of the dropout function (p_drop_) in the hidden layers was done (after every fully connected layer, the dropout layer was done). The L2 regularization term is also incorporated into the cross-entropy loss function. This regularisation term is controlled by the hyperparameter α, which determines the weight assigned to the regularisation. The network is trained via the Adam optimizer, employing a learning rate denoted as *lr*. The conclusion of the training process is regulated by an early stopping with a maximum of 2,000 epochs. In our model, DNN achieved an accuracy of 72% with AUROC = 0.85. **Table S4** gives the list of hyperparameters and selected values.

## 6. Discussion

It is often recognised that NSCLC has no symptoms and can only be found physically when it is very early on. Effective analysis and investigation of cancer genes to identify tumour biomarkers has been a popular issue for early cancer diagnosis expression patterns and treatment with the development of bioinformatics [50]. Overall, our machine learning classifier showed that the lung tissue microbiome-based binary-class model for subtype prediction is feasible for early screening of NSCLC. The novelty lies in the superior and reproducible machine learning and deep learning methods for classifying subtypes which is of high clinical relevance. We believe this binary-classification lung tissue microbiome-based model for distinguishing AC vs. SCC has the potential in clinical interventions and serves as a minimally-invasive way of early screening of this disease in clinical practice. Our results also have implications for the potential development of bacterial biomarkers for predicting subtypes using the identifien d specific markers for this disease. The strength of our findings includes two accurately classified NSCLC cohorts with AC and SCC with the help of lung tissue microbial profile.

Further, our machine learning classifier with taxonomic features, including phylum Deinococcota, order Enterobacteriales, genus Thermus, phylum Actinobacteriota, order Psuedomonales, genus Actinobacter, family Moraxelleace, and 11 others classified the two subtypes of NSCLC, namely AC, and SCC. Our study showed that lung tissue microbiome when complemented with state-of-the-art machine learning algorithms can discriminate the overlapped/closely related subtypes of NSCLC. This work has some limitations. Firstly, the disease spectrum of this study is still limited, and the inclusion of more classes can further enhance the value of this binary-class diagnostic tool. Secondly, while the models yielded high accuracy with fewer Type I and Type II errors, further study is required to validate in larger cohorts. Further studies should include a wider range of patients in terms of age, and geographic location as well as their medication and tumor markers to enhance the strength and applicability of the proposed model.

To the best of our knowledge, we present the novel lung tissue microbiome-based machine and deep learning binary classification model that achieved high performance for subtype classification. This microbiome-based model could potentially be applied clinically to complement early disease diagnostics and followed by monitoring of therapeutic interventions.

## 7. Conclusion

The study of metagenomics is gaining acceleration with advanced next-generation sequencing. This sequencing produces a feature table that taxonomically or functionally corresponds to a microbiome that, when correctly labeled can be used as inputs to ML and DL-based methods. These methods are a powerful tool with a wide variety of applications in the field of microbiome in LC. This study was the first to highlight the utility of ML and DL methods for binary classification using the lung tissue microbiome in NSCLC for early screening and better prognosis.

