## Supplemental Figure for "Machine learning analysis of lung adenocarcinoma and squamous cell carcinoma microbiome datasets reveals biomarkers for early diagnosis"

**Supplementary RP1**

**Figures S1:**
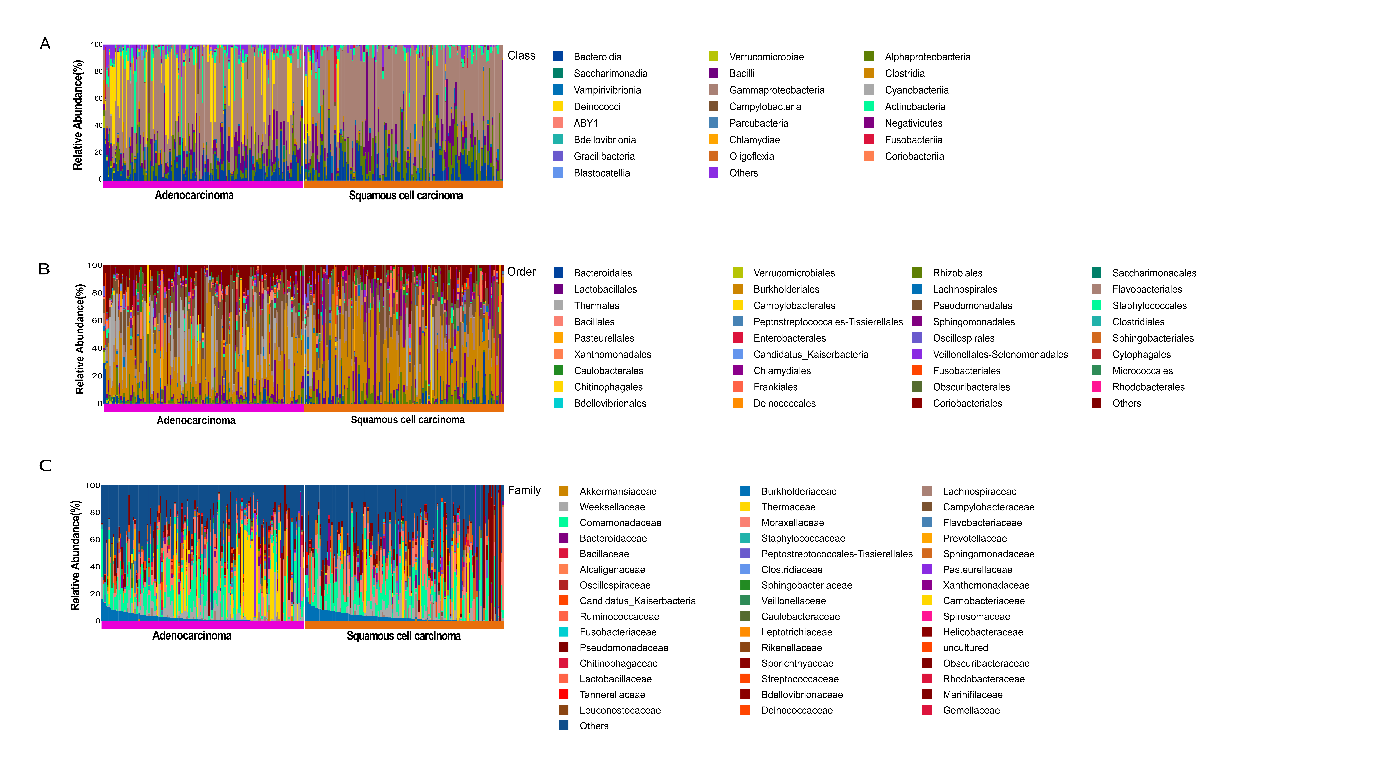

**Figures S2:**
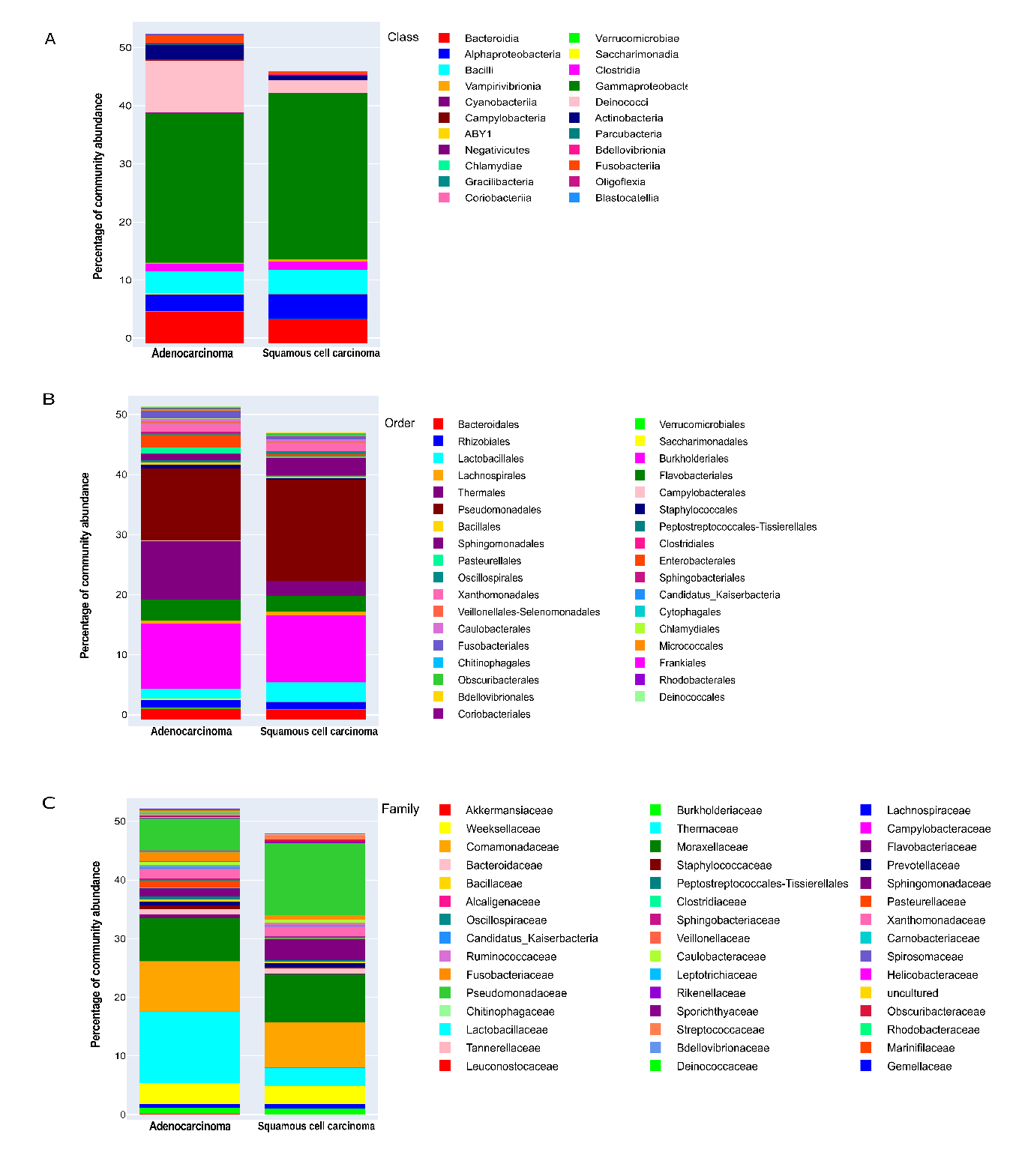

**Figure S3: Heatmap coorelation**

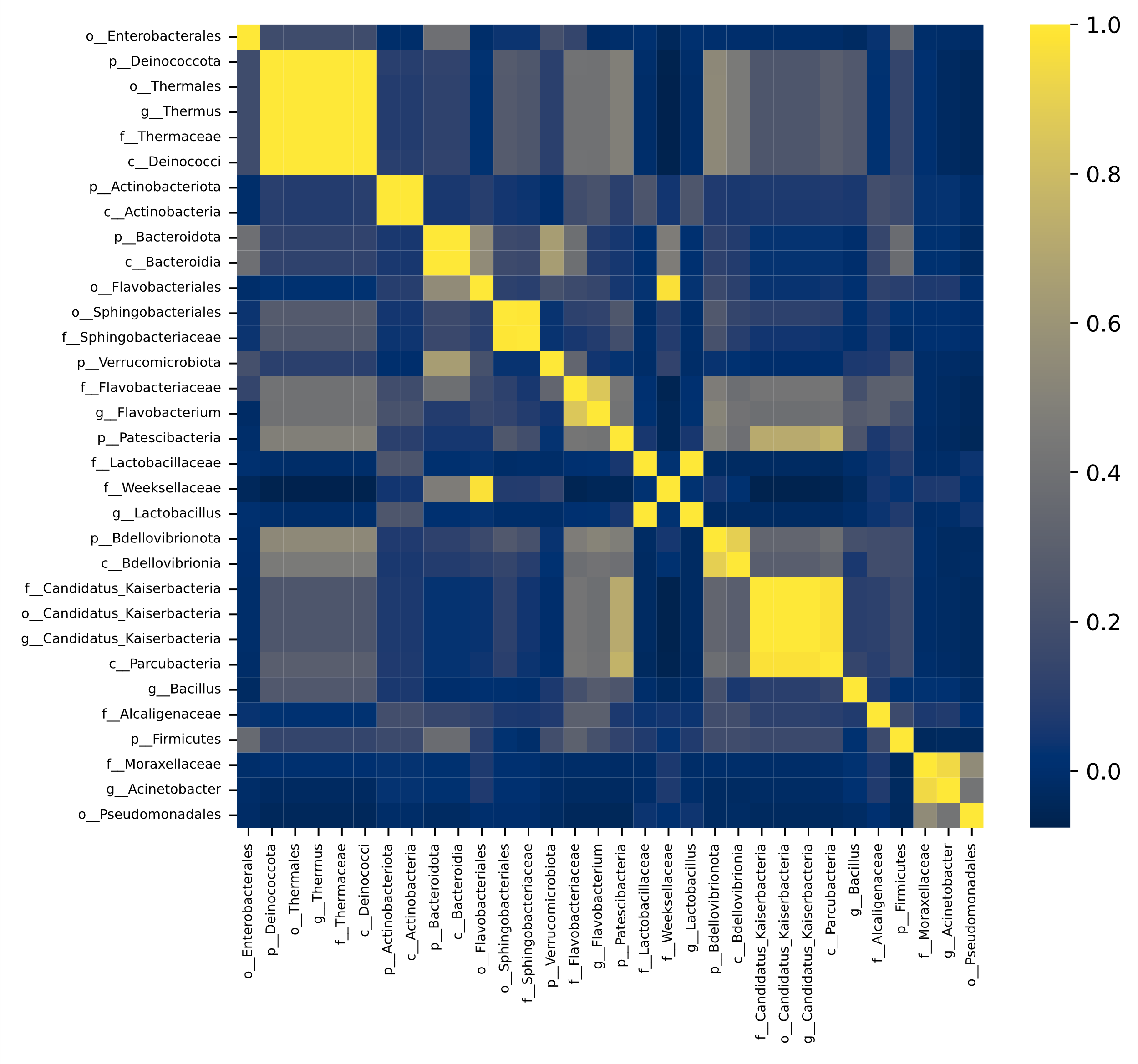

**Figure S4: Transforming dataset using LDA**

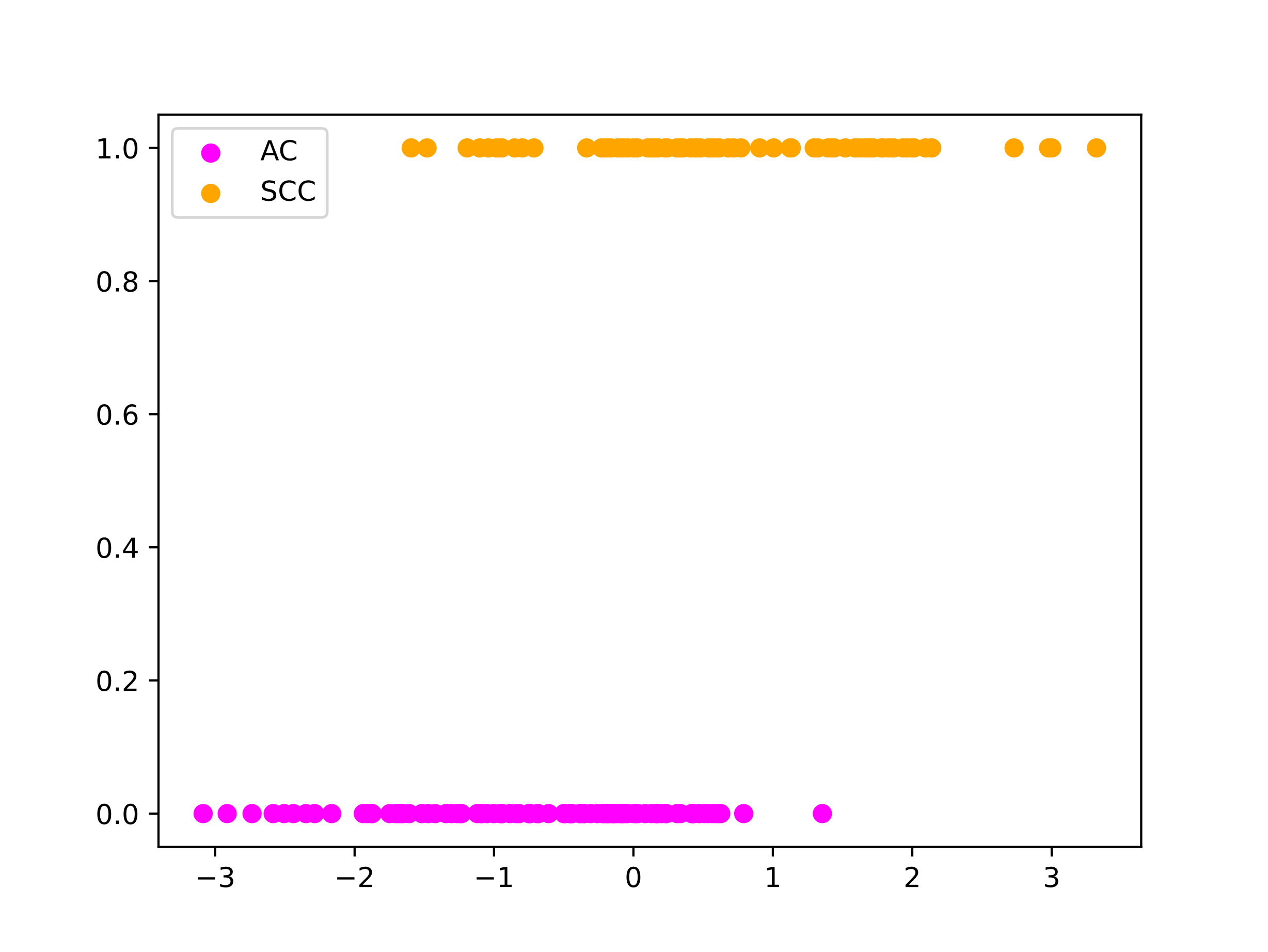
